# In-Silico Analysis of nsSNPs Associated with CYP11B2 Gene

**DOI:** 10.1101/602615

**Authors:** Anam Arooj, Muhammad Tariq Pervez, Zeeshan Gillani, Tahir Ali Chohan, M. Tayyab Chaudhry, Masroor Ellahi Babar, Asma Tufail Shah

**Author notes:** **Corresponding Authors: Dr. Muhammad Tariq Pervez**, Contact. **Dr. Zeeshan Gillani**, Contact.

## Abstract

*CYP11B2* gene is located over the upper layer of the kidney. It produces aldosterone synthase enzyme and thereby has an essential role to balance salt and mineral level in the body. A mutation in this gene can deregulate the production of aldosterone hormone in the body which may lead to many diseases including hypertension and cardiac diseases. To control the excess production of this aldosterone an inhibitor “*Fadrozole*” is being used which is associated with an active site cavity of CYP11B2. This study has been divided into two parts. In the first part, the four computational tools (SIFT, Polyphen-2, I-Mutant, ConSurf) were used to identify 29 deleterious SNPs out of 1600 *CYP11B2* SNPs. In the second part, five residues (R448G, R141P, W260R, F130S, and F445S) were identified in the active site cavity (out of 29 deleterious CYP11B2 SNPs) at the distance of 5A°. Binding free energy calculation as well as Dynamics simulation techniques were applied to determine the effect of these mutations on the CYP11B2-Fadrozole compound. The results showed that *Fadrozole* binding with CYP11B2 became stronger which proved the efficiency of this drug inhibitor with these highly damaging mutations. Our study will be useful for selecting the high priority CYP11B2 mutations, which could be further, investigated in this gene-associated study, for better understanding of the structural and functional aspects of the observed (CYP11B2) protein.

## 1. Introduction

Human genetic variations are critical in medical and functional perspectives. SNPs (Single Nucleotide Polymorphism) belongs to most abundant classes of genetic variations in the human genome. These contributes in many complicated central nervous system phenotypes, including drug response, susceptibility to neuro-physiological as well as psychiatric disorders. These mutations are highly abundant with a frequency of 1 to 4 out of every 1,00 bases in the human genome [1]]. SNPs are of different types, mostly they are neutral but some of them are functional that can cause an amino acid modification which ultimately can affect protein structure and function, these are known as missense SNPs / nsSNPs (non-synonymous SNP).

Scientists have been trying to identify the functional SNPs of human genes for different diseases [2]]. Although lab experiments can accurately identify the genetic role of an SNP [3] but it is not practically possible to test a large number of genetic variants in lab even for a single gene. For example, a single gene like **CYP11B2** has more than 1600 SNPs (at the time writing article) which raises a need to prioritize SNPs to minimize this big number to a practical number for lab experiments [4]. This necessity leads researchers to computational study of the genetic variants to predict disease-associated amino acid mutations. The computational tools are fast and reliable in their predictions with accuracy of 80-85% [5].

CYP11B2 is located over the upper layer of the kidney at position 8q24.3 [5]]. It produces aldosterone synthase enzyme and thereby has an essential role to balance salt and mineral level in the body. Any SNP in this gene can deregulate the production of aldosterone hormone in the body which may lead to many diseases including hypertension and cardio vascular disease. [6]]. Brand E, et. Al., found a positive association of CYP11B2-344T allele with essential hypertension [8–9]. During a study, 175 cardiac patients of European Continental Ancestry Population were diagonosed and it was identified that C allele (CT, CC) at −344T/C SNP position in “aldosterone synthase” gene does not considerably effect clinical prognosis of cardiac heart-failure [7]. Similarly the excess of aldosterone has been reported for hypertension [8] which is one of the most prevalent diseases these days around the world and affects more than 1 billion people worldwide[12]. Regular hypertension control medicines work fine but still a large number of patients don’t get benefit of them and their blood pressure remains high even after treatment [13]. One of the possible solutions used by scientists is the use of an inhibitor (Fadrozole) to limit the biosynthesis of aldosterone hormone [14].

Jia et.al [9] studied CYP11B2 gene for 52 SNPs and found four SNPs as deleterious. The novelty of this study was that we filtered out 29 deleterious SNPs from a large number of non-risk alleles and studied the effect of Fadrozole drug over some pathogenic SNPs. There are more than 1600 identified SNPs in CYP11B2 gene but only one mutation (−344T/C rs1799998) got primary focus for disease association studies. In this study, we performed computational analysis of CYP11B2 gene to identify the pathogenic mutations. The second objective of this study was to identify the effect of missense mutations over receptor-ligand (CYP11B2-Fadrozole) binding; whether these mutations strengthen or weaken the inhibitor affinity with CYP11B2. Fadrozole is an inhibitor used to limit the excess production of aldosterone from CYP11B2 to control diseases e.g. cardiovascular and hypertension. Aldosterone synthase responsibility is to biosynthesize the main mineralocorticoid aldosterone in the adrenal cortex.

A computational study on the same gene was performed in 2014 [15] but we monitored four times increase in SNPs (from 358 to >1600: in 2017 when we thought to study this gene) which motivated us to re-investigate nsSNPs associated with CYP11B2 gene. Our study will help biologists to further understand the CYP11B2 gene association with human diseases, and for future experiments.

## 2. Materials and Methods

### 2.1 Dataset collection

The SNP information i.e. SNPID, Gene Position, mRNA position, Protein Position, Residue Change, mRNA (accession number: NM_000498.3), Protein Sequence (NCBI Accession number: NP_000489.3) of the human CYP11B2 gene were obtained from NCBI (National Center for Biotechnology Information) database of SNPs (dbSNP (http://www.ncbi.nlm.nih.gov/snp/) [10]. The crystal structure of CYP11B2 protein in complex with inhibitor Fadrozole (PDB ID: 4FDH) were obtained from RCSB protein Data Bank (www.rcsb.org). In this study, these (ligands binding) regions were used as active sites for docking. Figure 1 shows the workflow for in-silico analysis of functional SNPs in the human CYP11B2 gene.

**Figure 1.**
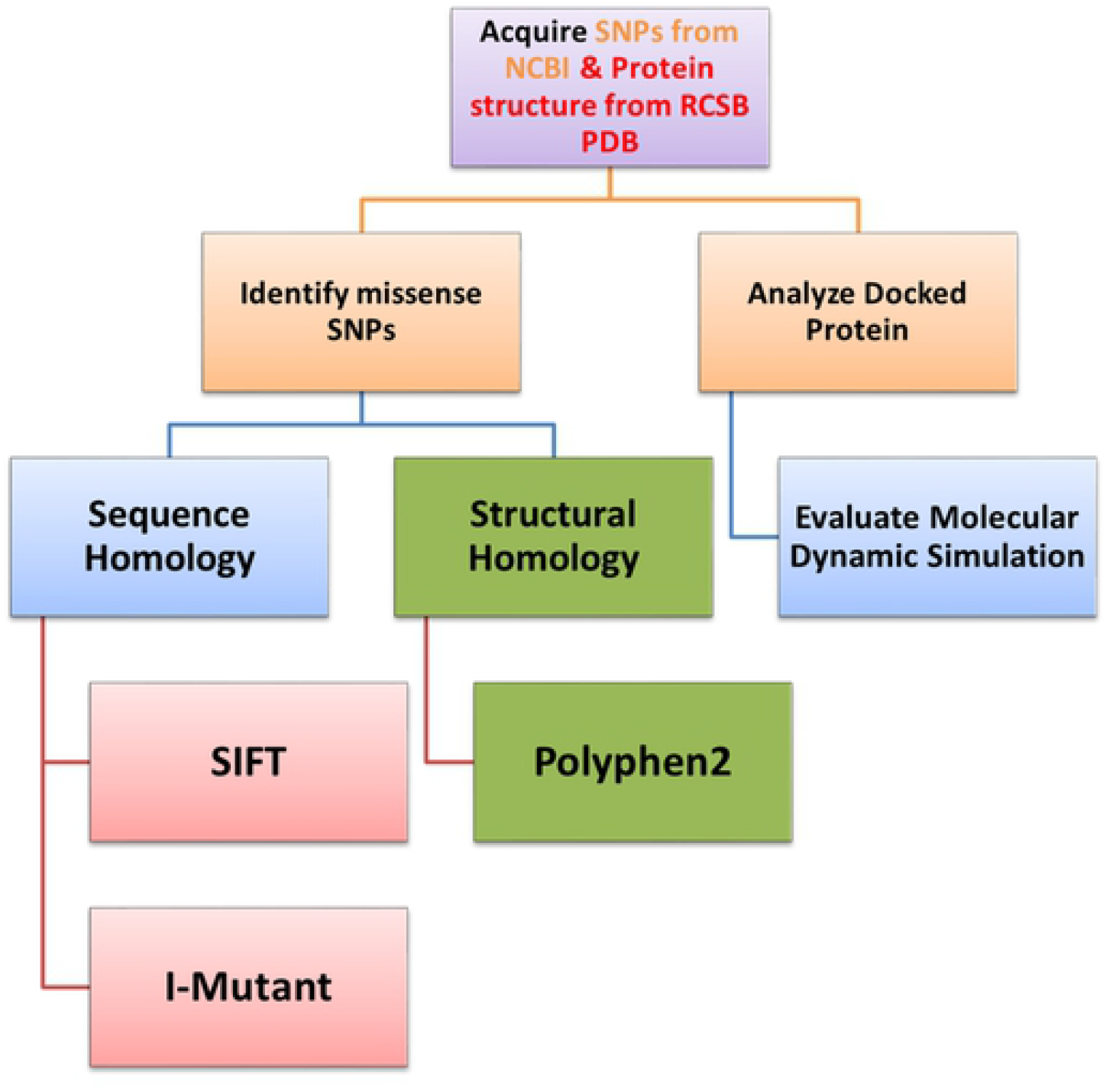
shows the workflow, tools, and databases used for in-silico analysis of functional SNPs in the human CYP11B2.

### 2.2 Bioinformatics Tools used to identify deleterious SNPs

In order to evaluate potential impact of the selected missense SNPs, we utilized four prediction tools and filtered all missense SNPs that were classified as deleterious by all of them. These computational tools include SIFT, I-Mutant, Consurf and Polyphen-2.

*SIFT (Sorting Intolerant From Tolerant)* is a sequence homology based bioinformatics tool which can be accessed at (http://sift.bii.a-star.edu.sg/index.html) [11]. This tool was used to predict whether the protein will tolerate the newly introduced amino acid caused by SNP mutation or it will be proved as a damaging mutation [16]. Genome tool “Sift non-synonymous single nucleotide variant (human build 37)” was used in our study. The SIFT input query string includes Chromosome number (8 for CYP11B2), Chromosome position, Orientation and allele respectively. SIFT outputs tolerance index score (Score), SNP mutation of index score < 0.05 are considered as deleterious. A very low tolerance index indicates that the particular mutation has a more functional impact [12]].

*Polyphen-2* is a protein sequence and structure based method. This tool was used to foresee the amino acid substitution impact on the structure and function of human protein. This can be accessed at (http://genetics.bwh.harvard.edu/pph2/)[13]. Polyphen-2 query was performed by submitting SNP Identifier/Accession #, protein position and Substitutions / Residue changes. This computational tool output a large heat-map color bar that range from 0 to 1 to show damaging probability of an SNP. A high score shows the higher chances of a mutation as damaging.

The protein stability over single mutation was measured using *I-Mutant Suite which can be* accessed at (http://gpcr2.biocomp.unibo.it/cgi/predictors/I-Mutant3.0/I-Mutant3.0.cgi) [14]. Its input includes protein position, new residue information and protein sequence (obtained from Uniprot against entry P19099). The output was used to classify Predicted Free Energy change value (DDG) of protein. DDG < −0.5 (kcal mol^-1^) shows huge decrease in protein-stability on SNP mutation which is critical.

Highly conserved regions in proteins/nucleic-acid-sequence often indicate structural and/or functional importance [18]. Consurf web server which can be accessed at (http://consurf.tau.ac.il/2016/)[15]. It calculates the evolutionary conserved amino acid positions in proteins using Empirical Bayesian inference (protein structure and sequence respectively). Amino acid analysis was performed for PDBID (4FDH) and identifier ‘A’. The conservation scores were calculated among protein and its homology whose color scale ranges from 1 to 9, where the scores in the range 7 to 9 indicate conserved region, probability in ascending order.

### 2.3. Molecular Docking

The co-crystallized structure for CYP11B2 bonded to Fadrozole (CYP11B2-Fadrozole complex) was retrieved from the Protein Data Bank (PDBID: 4FDH). The initial conformation of Fadrozole in 4FDH has been used as starting point for molecular docking. Prior to Dock the inhibitor into the active site, the receptor-protein structure was prepared (with the help of structure preparation tools) to validate the chemical accuracy in biopolymer module of SYBYL-X 1.3[16]. For that purpose, at first step missing hydrogen atoms added, then all crystallographic waters molecules were removed, then atom-types were assigned, next, atomic charges were applied according to AMBER 7 FF99 force-field. The Power algorithm was used to minimize the energy of each complex, for 1000 cycles at convergence-gradient of 0.5 kcal (mol A°). The backbone atoms were kept fixed during structure optimization.

Five different in-silico mutants of CYP11B2 (R448G, F445S, W260R, R141P and F130S) were generated by replacing Arg448, Phe445, Trp260, Arg141 and Phe130 with Glycine448, Serine445, Arginine260, Proline141 and Serine130 amino acids respectively. All mutations were induced using SYBYL-X 1.3 (Biopolymer module)[16]. Prior to molecular docking, the generated mutant models of CYP11B2 were energy-minimized and MD-simulated at least for 10 ns, which have been discussed in a later section. For molecular docking, the active structural conformation of Fadrozole was extracted from its complex 4FDH.

Finally the active sites of these five mutants of CYP11B2 were docked by the ligand using the same protocol as reported in our previous publications [10–12]. Twenty best-docked conformations were retrieved for each ligand-mutant complex system. The putative ligands poses were ranked according to Hammerhead scoring function (C-score) [4]. To examine the effect of the point CYP11B2 mutation over receptor–ligand interaction, all ligand protein complexes were visualized and critically investigated.

### 2.4 Molecular dynamics Simulation

For all five generated structures of the complexes (Fadrozole-CYP11B2, Fadrozole-R448G, Fadrozole-F445S, Fadrozole-W260R, Fadrozole-R141P and Fadrozole-F130S) was further stabilized by performing MD simulations. These simulations were performed with the SANDER module in AMBER12 software package in a solvation system. LEaP program embedded in AMBER12 was used to add missing hydrogen atoms of protein. The complex was immersed into an octahedron box of TIP3P water molecules to achieve water molecules up-to minimum 10°A distance between protein and edge of box. Extra water molecules were replaced by Cl^-^/Na^+^ counter-ions by using LEaP protocol.

AMBER ff99SB force field, for protein, and General AMBER force field (GAFF together with RESP charges), for ligand, was used to assign force field parameters for each protein-ligand complex.

Each protein-ligand complex of the mutant and wild type was then subjected to production simulations run of 50 ns by keeping protocol and parameters same as those reported in our previous publications [10–12]. All these analyses were performed using ANAL, CARNAL and PTRAJ modules of ‘AMBER12’ software.

### 2.5 MM/PB(GB)SA Based Free energy calculation

MM/PB(GB)SA [13], a molecular-mechanics based scoring method, was used to compare the binding free energies of Fadrozole-R448G, Fadrozole-F445S, Fadrozole-W260R, Fadrozole-R141P and Fadrozole-F130S mutant complexes to wild type Fadrozole-CYP11B2. The pairwise nature of the GB methodology allows to produce insightful interaction and desolation components by decomposing free-energies. All MM/PB(GB)SA calculations [13] were thoroughly performed in the AMBER16 software package, while keeping protocol and parameters same as provided in our previous publications [10,12].

## 3. Results

### 3.1 Identification of Deleterious nsSNPS

The in-silico analysis for 275 missense SNPs of CYP11B2 (These were total nsSNPs when we started work; now there is more than 600 nsSNPs) was performed (Table S1: Given in Supporting Information Table S1) using SIFT, I-Mutant, Polyphen-2 and computational bioinformatics tools. SIFT was run for 275 SNP (Table S2) and it predicted 98 (35.63%) nsSNPs as damaging, while 162 were of benign effect. Polyphen-2 was run for 275 SNP out of which 128 SNPs (46.54%) were predicted as damaging (Table S3) while 147 mutations were resulted as benign; I-Mutant (structure based tool) was run for total 186 nsSNPs (67.63%) which resulted that 117 nsSNPs (62.9%) show a DDG < −0.5 hence they are largely unstable (Table S4).

The combined results of these three computational tools screened down the CYP11B2 nsSNPs from 275 to 57 (20.72%). Thus 57 mutations (Given in supporting information S5 Table) were predicted by all these three tools that had higher probability of damaging protein structure and cause disease. Figure 2 shows the distribution of benign and deleterious and nsSNPs found using I-Mutant suite, Polyphen and SIFT.

**Fig 2.**
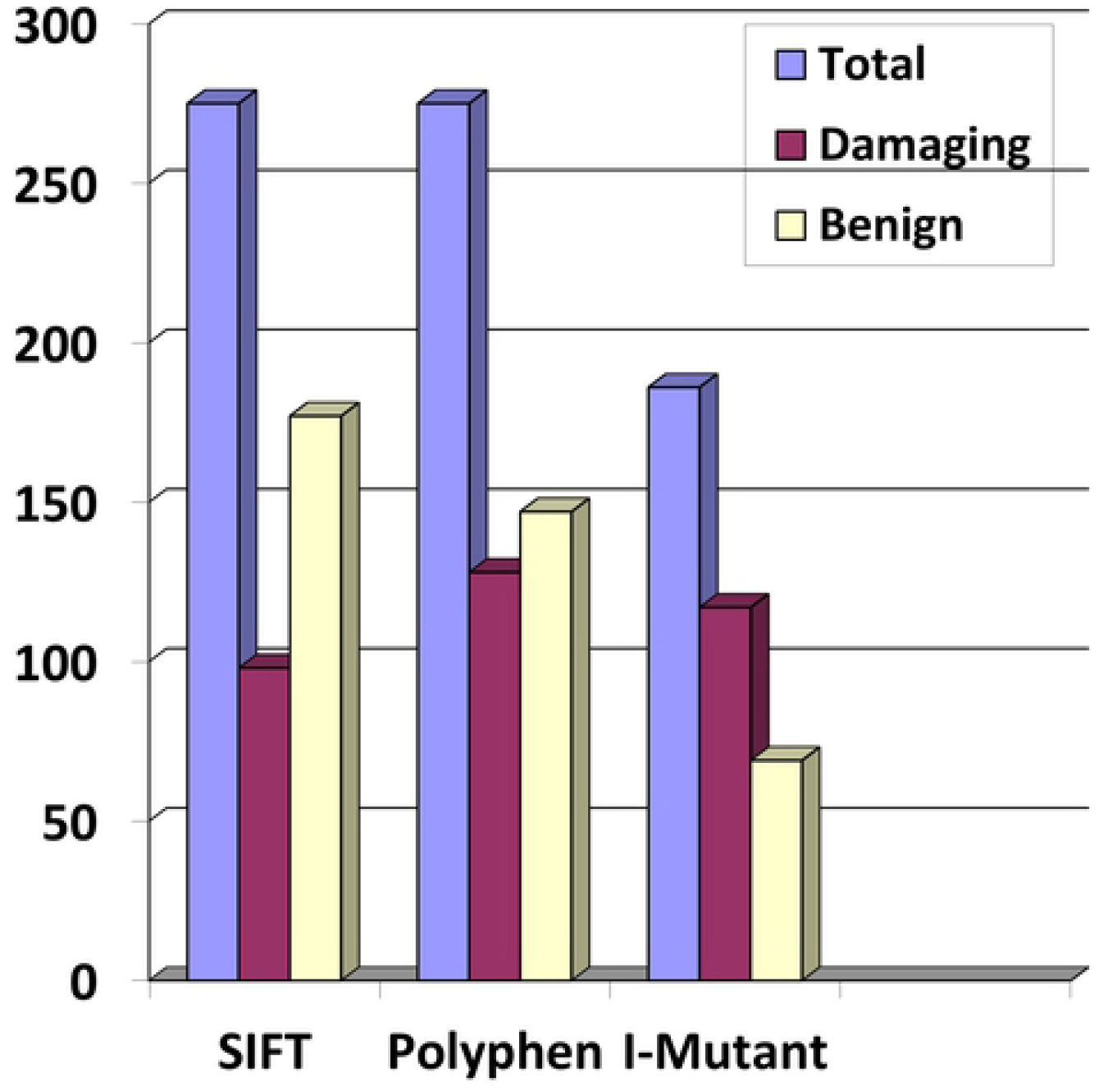
Distribution of deleterious and benign nsSNPs by bioinformatics tools Polyphen-2, SIFT, and I-Mutant Suit.

### 3.2 Evolutionary Conservation analysis of nsSNPs

It is a known fact that mutation in highly conserved protein regions keeps more chances to cause disease against any mutation. The 4FDH.pdb protein structure was analyzed using Consurf to find the conserved regions. ConSurf identified 174 conserved amino acid positions of 4FDH (Table S6). When compared with 57 nsSNP (predicted as damaging by Polyphen, I-Mutant and SIFT), only 29 nsSNPs were found in conserved area (Table 1).

**Table 1:**
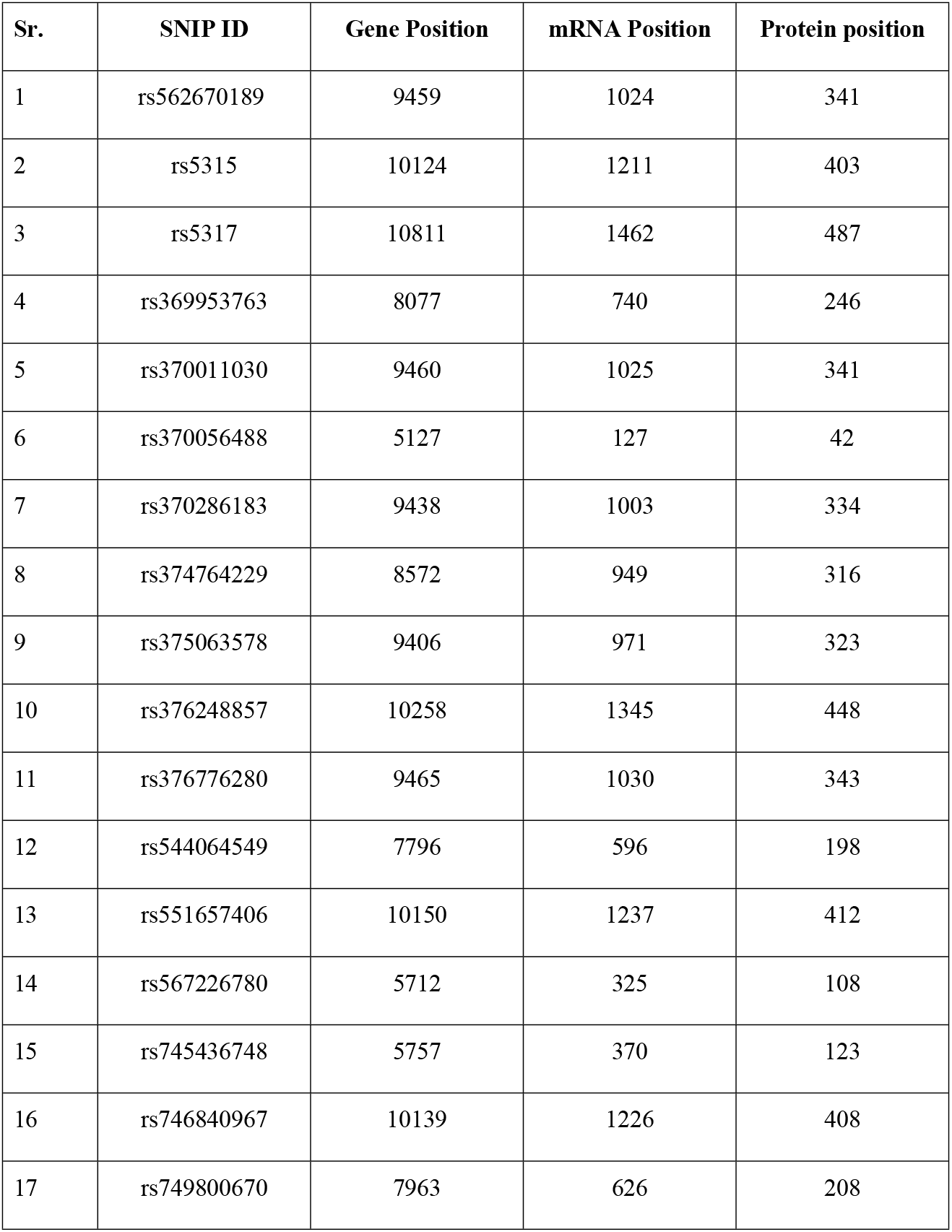
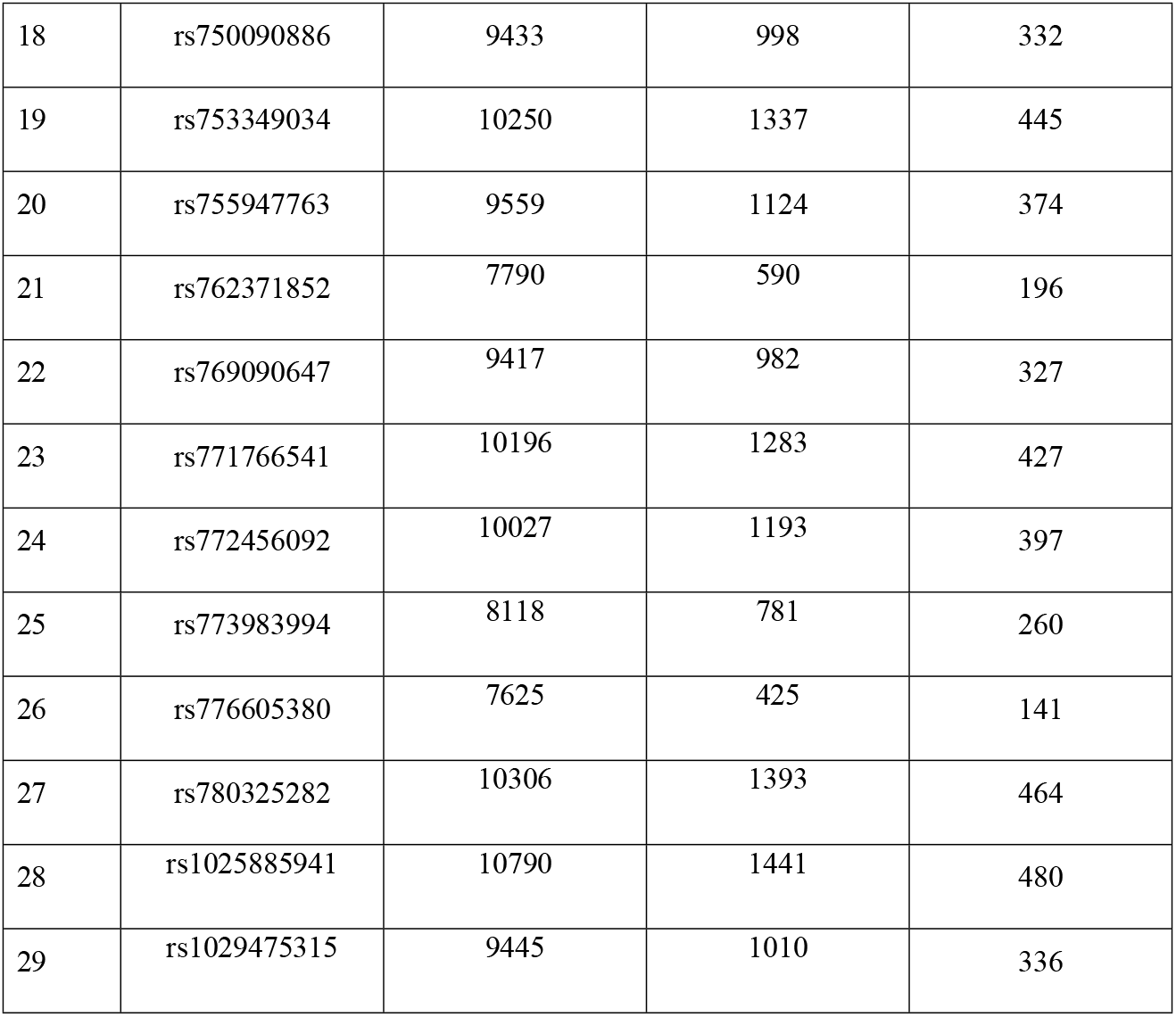
Damaging nsSNPs by SIFT, POLYPHEN, I-Mutant and ConSurf

### 3.3 Molecular Docking

The docking identified 18 residues at the distance of 5Ȧ from the active site of CYP11B2-Fadrozole compound, these were R110,W116,R120,F 130,W137,R141,F231,W260R, A313, S315, T318, L319, L373, A384, G445, R448 and C450. The next step was to filter out only those residues out of 18 residues which were identified as damaging (Table 1), hence only five residues were subjected for further analysis. R448G and R141P reside were found at 3 Ȧ from drug fadrozole in CYP11B2-fadrozole complex (Fig 3A) while G445, W260R and F130S reside at the distance of 5 Ȧ shown in (Fig 3B).

**Fig 3.**
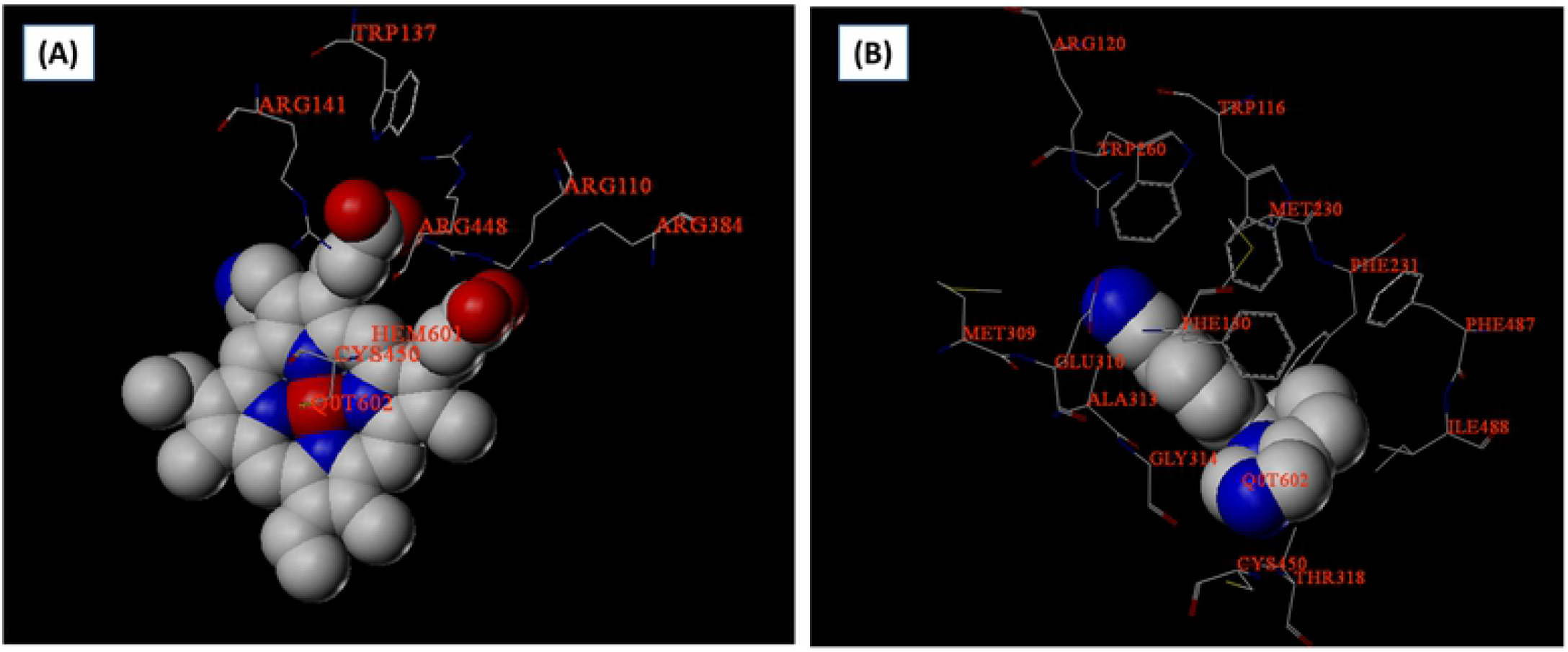
Amino acids at different distances from drug Fadrazole (A) at 3 Ȧ distance from drug Fadrozole, (B) at 5 Ȧ distance from drug Fadrozole

**Table 2:** Damaging Conserve Area SNPs near Fadrozole-CYP11B2 compound

| Sr. | SNIP ID | Amino Acid change | Protein Position |
| --- | --- | --- | --- |
| 1 | rs376248857 | R448G | 448 |
| 2 | rs753349034 | F445S | 445 |
| 3 | rs773983994 | W260R | 260 |
| 4 | rs776605380 | R141P | 141 |
| 5 | rs965343088 | F130S | 130 |

Docking results revealed that there were no particular changes in binding modes of ligands in mutant and wild-type, however, this was not surprising because the binding pocket was almost the same and only one residue was different. The docking scores revealed that the ligand bonded to the SNPs with higher affinity than wild-type/native; however, to further confirm the docking results molecular dynamic simulations were initiated.

### 3.4 Molecular Dynamic Simulation

Although, molecular docking analysis is sufficient to depict ligand-receptor interactions. The docking can be further refined with more reliable molecular dynamic (MD) simulation approach. As docking scores cannot be considered as 100% reliable to distinguish ligands on their selectivity basis. Moreover, in molecular docking approach physiological conditions are not considered. Thus, post-processing molecular docking results with MD simulation may would be helpful to identify dynamic stability of ligand-protein complex. To clarify the structural consequences due to these five identified mutations, docking results were subjected to Molecular Dynamic (MD) simulation analysis.

To examine the stability of the Fadrozol-CYP11B2 complex stability under the simulation conditions the RMSD (root mean square deviation) of the trajectories was measures from their initial structure. The analysis of the RMSD for the active site residues around 5 Ȧ of ligand backbone atoms and ligand heavy atoms performed for 40ns are observed.

Figure 4 shows that the protein backbone atoms RMSD of Fadrozole inhibitor bound CYP11B2 mutant complexes (F130S, F445S, R141P, W260R and R448G) in the simulation fluctuates around 2.2 A°, 1.6 A°, 1.8°A, 2.3°A and 1.7°A after 10000 ns, 15000 ns, 25000ns,18000ns, and 15000 ns, respectively. These results reveals the stability of the docked complex structure throughout MD simulation, although protein structure shows some fluctuations, while the ligand exhibits smaller conformational changes. The low fluctuation in RMSD plot demonstrate that the results are in accordance with ligand-protein (residues) high affinity that are forming a stable complex. Among five ligand-receptor complexes F445S exhibited lowest atomic fluctuations. Hence, it concludes the ligand stability in the binding pocket such that it did not dissociate from the protein during the simulation. Overall, these analyses for ligand-protein complex stabilization suggest that the simulated docking conformation are correct and can be used to calculate binding free energies.

**Fig. 4.**
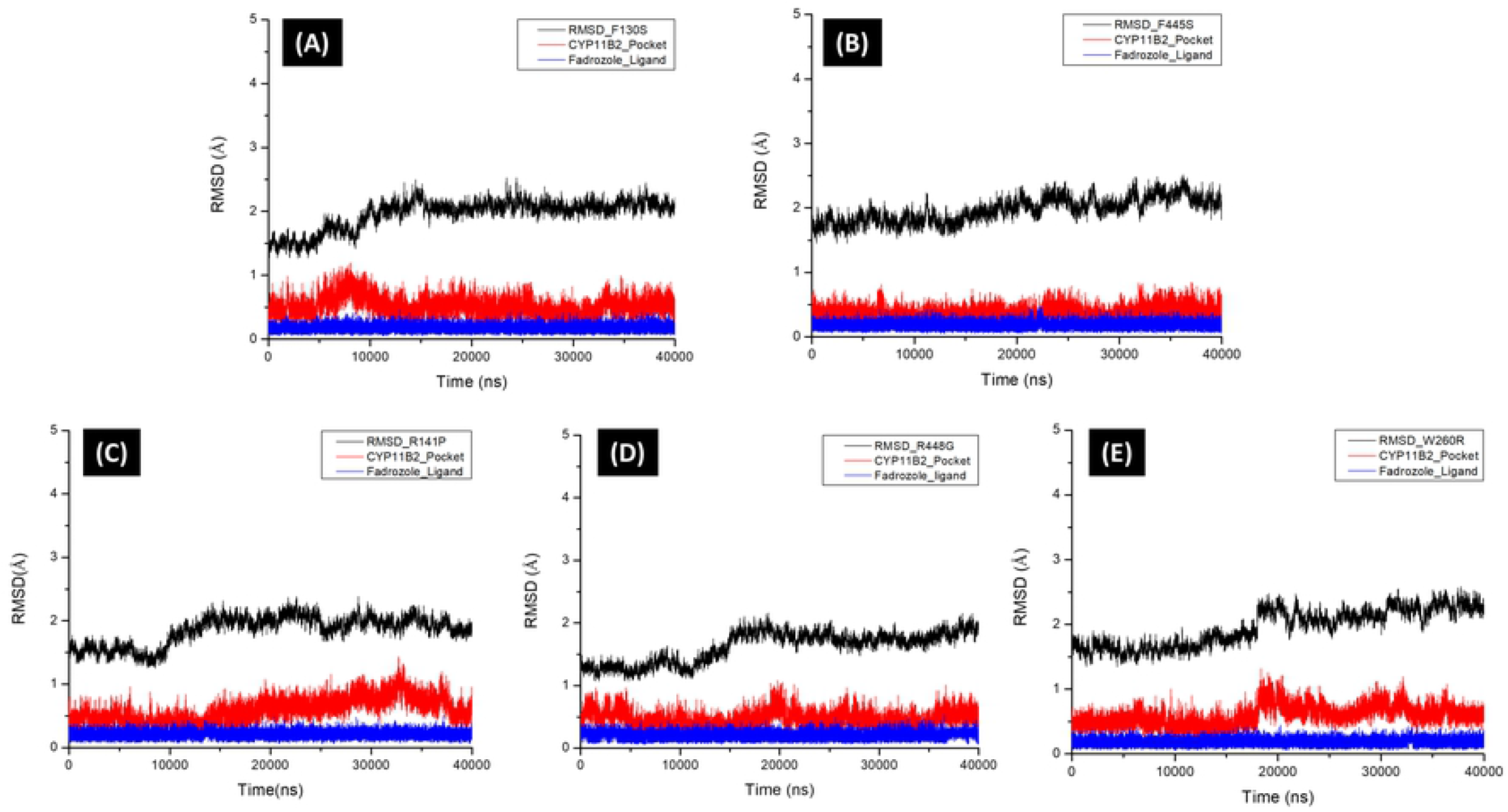
The RMSD of the backbone atoms of protein (black), backbone atoms of binding pocket residues around 5 °A of ligand (red), and the heavy atoms in the ligand (blue) for (A) F130S (B) F445S (C) R141P (D) R448G and (E) W260R bound CYP11B2 in complexes with respect to the initial structures as a function of time 40ns (nanosecond).

#### 3.4.1 Binding free energy analysis

The binding affinity of ligand towards CYP11B2 and its mutations were calculated using Molecular Mechanics/Poisson-Boltzmann Surface Area (MM/PBSA and MM/GBSA) methodologies[17]. The binding free energy (BFE) was calculated in terms of solvation energy and entropic contributions over 1000 snapshots for the complex, receptor and ligand respectively, obtained over last 5 nano-seconds of MD trajectories for protein-ligand system. Table 3 and Fig 3 (A-E) summarizes computed BFEs and their enthalpy and entropic contributions for five systems (fadrozole-R448G, fadrozole-R141P, fadrozole-W260R, fadrozole-F130S and fadrozole-F445S). The Fadrozole inhibitor possesses higher predicted binding energies for F445S (ΔG_pred_(GB) MMGBSA: −19.88 kcalmoli^-1^; MMPBSA: −28.13 kcalmol^-1^) and indicate more significant difference in ΔGpred(_GB/PB_) values for F445S vs R448G, R141P, W260R and F130S. Still it can be seen that the binding energy increases for all five mutants of fadrozol-CYP11B2 than wild type.

**Table 3:** Binding free energy change due to five mutations in the Fadrozole-CYP11B2 compound.

| **Protein-inhibitor** | **Wild-Type**<br>**(4FDH)** | **4FDH_R448G** | **R141P** | **W260R** | **F130S** | **F445S** |
| --- | --- | --- | --- | --- | --- | --- |
| $\Delta E_{vdW}$ | -24.98 | -27.35 | -33.18 | -29.59 | -29.12 | -29.58 |
| $\Delta E_{ele}$ | -15.39 | -5.36 | -10.34 | -2.16 | -10.50 | -22.77 |
| $\Delta G_{nonpol, sol}$ | -3.07 | -3.79 | -4.41 | -3.96 | -3.71 | -3.91 |
| $\Delta G_{ele, sol} (PB)$ | 30.89 | 21.95 | 27.78 | 17.84 | 29.82 | 35.56 |
| $\Delta G_{ele, sol} (GB)$ | 23.76 | 14.43 | 17.76 | 10.34 | 19.40 | 28.11 |
| $\Delta E_{vdW} + \Delta G_{nonpol, sol}$ | -28.05 | -31.13 | -37.59 | -33.55 | -32.83 | -33.48 |
| $\Delta E_{ele} + \Delta G_{ele, sol} (PB)$ | 15.50 | 16.59 | 17.43 | 15.68 | 19.32 | 12.80 |
| $\Delta E_{ele} + \Delta G_{ele, sol} (GB)$ | 8.37 | 9.07 | 7.42 | 8.18 | 8.90 | 5.35 |
| $\Delta G_{pred} (PB)$ | -12.60 | -13.99 | -19.05 | -17.27 | -12.84 | -19.88 |
| $\Delta G_{pred} (GB)$ | -19.68 | -22.06 | -30.17 | -25.38 | -23.93 | -28.13 |

In order to identify the driving forces for selective bindings of ligand on inhibitor, total binding free energy was decomposed into independent (binding free energy) components. (Fig 34-E or Table 3) using MMGB(PB)SA methods.

The calculated values of individual binding free energy components for four systems (Fig. 5 A-D) revealed that the disfavor-able electrostatic interactions between protein and ligand (**ΔE_ele_**) in vacuum were opposed by the favorable electrostatic energy of solvation (**ΔG_ele,sol_**). However, the sum of the electrostatic interaction contributions of (**ΔE_ele_**) and (**ΔG_ele,sol_**) slightly disfavors the protein-ligand binding. F445S shows the opposite behavior, (**ΔE_ele_**) decreases and (**ΔG_ele,sol_**) increases, and sum of these two favors this binding. The **sum of** vdW energy (**ΔE_vdW_**) and nonpolar solvation energy (**ΔG_nonpol,sol_**) is favorable contribution to inhibitor binding to CYP11B2 for all five mutant systems. The total vdW contribution for fadrozole-CYP11B2 complex was −28.05 kcal.mol^-1^. However, total predicted binding free energy in terms of both **ΔG_pred (PB)_** and **ΔG_pred_ (_GB_)** was reduced for all mutants and maximum favorable binding was observed for R141P (−19.05 kcal.mol-1) and F445S (−19.88 kcal.mol) nsSNPs mutants.

**Fig 5.**
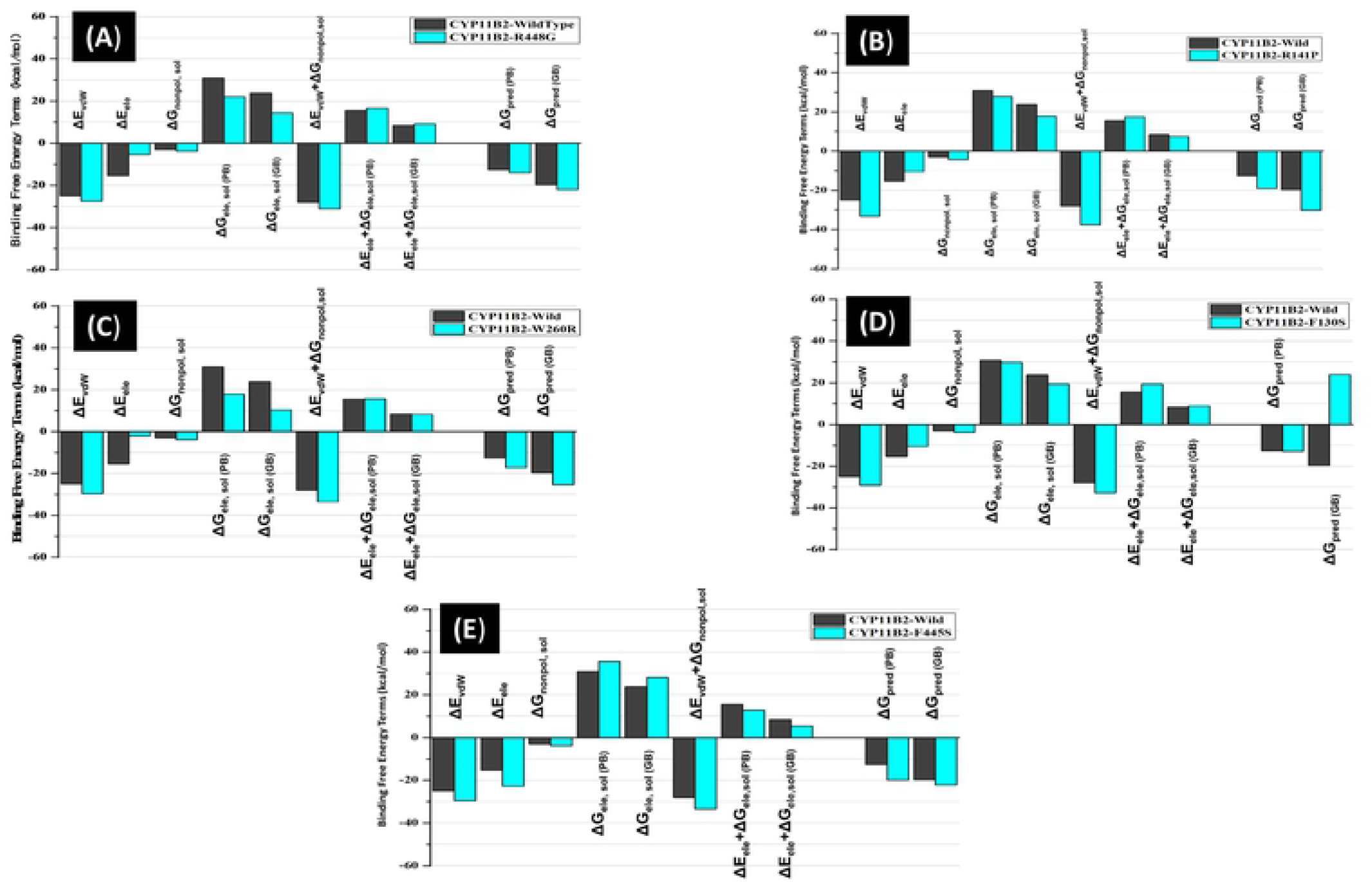
Comparison between binding free energy terms of Fadrozole (A) R448G, (B) R141P, (C) W260R, (D) F130S, (E)F445S

#### 3.4.2 Insight into the binding affinity of Fadrozole compound to CYP11B2

For all receptor-ligand complexes, the binding free energies were decomposed on interaction energies of Fadrozol with residues to estimate the role of distinct residue in tight binding of inhibitor to active site of protein. In case of R141, ten amino acids TRP83, ARG87, MET197, PHE198, GLU277, ALA280, GLY281, SER282, ILE449, GLU277 have favorable binding free energy, whereas five amino acids were disfavor-able (shown in Fig 6A-E). For F445S, mutant has low binding free energy for TRP83, ARG87, GLU277, ALA280, GLY281, PHE97, LEU98, LEU412 which contributed towards strong interaction with Fadrazole inhibitor.

**Fig 6:**
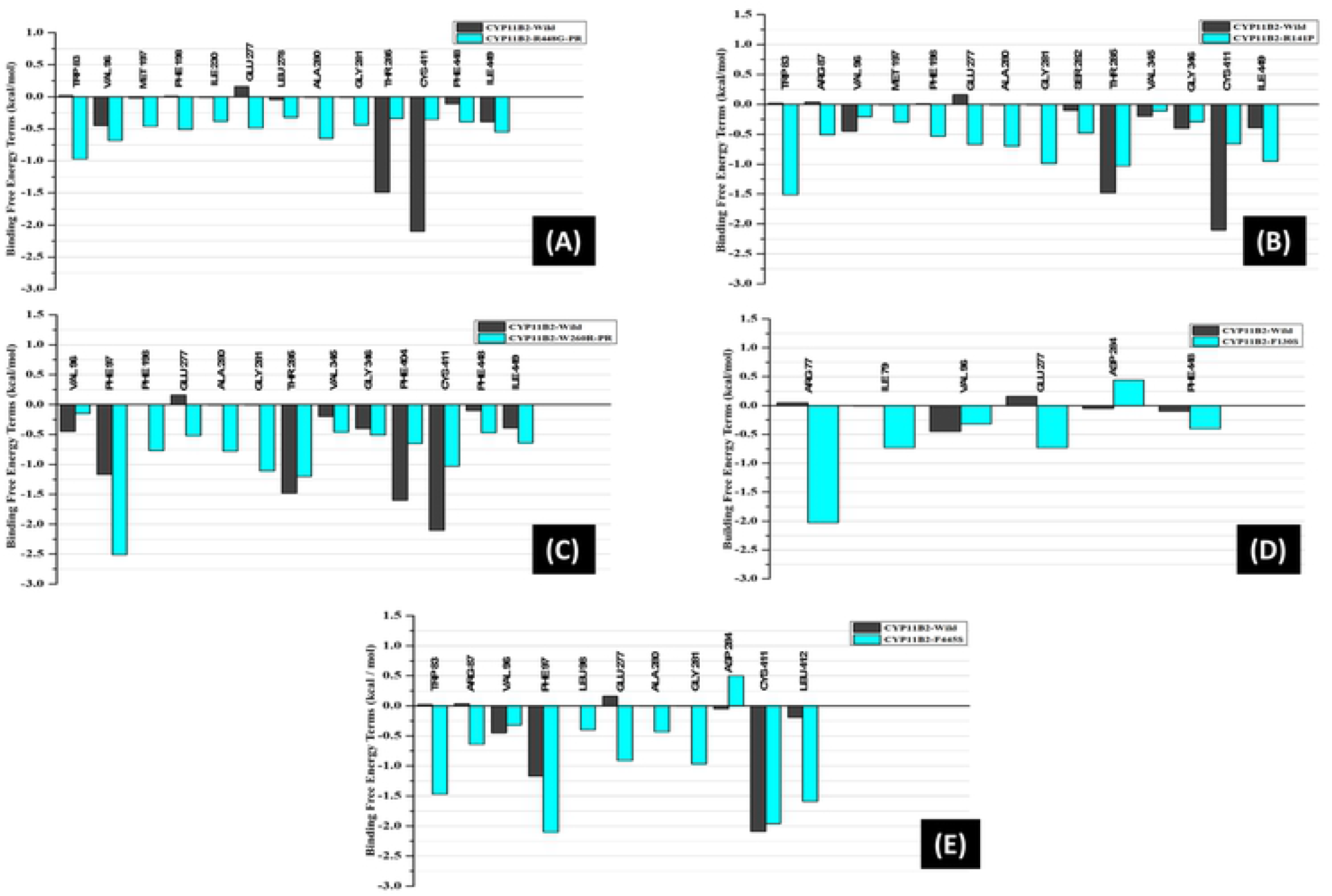
Comparison between energy contributions of residues around the binding site of CYP11B2-Fadrozole (A) R448G, (B) R141P, (C) W260R, (D) F130S, (E) F445S

Analysis of free energy components showed that increase in selectivity was directly associated with reduction of binding free energy of amino acids.

## 4. Discussion

The excess of aldosterone has been reported to cause the hypertension [8], which is one of the most prevalent diseases these days around the world and affected more than 1 billion people worldwide[12]. Regular hypertension control medicines work fine but still a large number of patients don’t get benefit of them and their blood pressure remains high even after treatment [13]. One of the possible solutions used by scientists is the use of an inhibitor (Fadrozole) to limit the biosynthesis of aldosterone hormone [14]. The inhibitor binds with CYP11B2 gene at a specific binding cavity. This binding strength between inhibitor and CYP11B2 gene depends on interaction energies of inhibitor with the residues of protein (active cavity site/ nucleotides). Any mutation in these nucleotides at binding sites may affect the binding free energies of receptor-ligand complex and hence can alter inhibitor efficiency. Almost 1600 SNPs of CYP11B2 has been reported till 2018 (https://www.ncbi.nlm.nih.gov/snp). In this study, we performed computational analysis of CYP11B2 gene to identify the pathogenic mutations and the effect of missense mutations over receptor-ligand (CYP11B2-Fadrozole) binding; whether these mutations strengthen or weaken the inhibitor affinity with CYP11B2 [18].

This study prioritizes SNPs with functional significance from an enormous number of non-risk alleles and provides new insights for further genetic association studies. Of 1600 SNPs, we selected about 275 **missense** SNPs for our investigations. In the first phase of our research we identified 57 deleterious nsSNPs using Sequence and structure based tools namely SIFT, Polyphen and I-Mutant (Table S7: Supporting Table S7), in second phase we identified evolutionary conserved areas of 4FDH (PDB of CYP11B2) (Table S6) due to the importance of the fact that mutation in evolutionary conserved areas are more liable for disease causing[19] and filter 29 of 57 highly deleterious SNPs that resides in conserved protein areas. In the third phase we add another dimension into our research, and check that if any of these nsSNPs has effect on drug Fadrozole. Docking and Molecular dynamic simulation studies have been performed to calculate the binding energy of Fadrozole-CYP11B2 complex. The 4FDH protein structure was analyzed in a docking software and it was found that five (R448G, R141P, W260R, F130S, F445S) out of 29 SNPs were located near fadrozole-CYP11B2 biding cavity. To further understand the structural consequences of the five identified mutations, docking results were analyzed by Molecular Dynamic (MD) simulation. The simulations were performed for both wild-type and mutant nsSNPs the results showed that binding energy of complex increased even after these deleterious mutations.

The literature on CYP11B2 SNPs with disease causing probability is scarce. One mutation (−344T/C rs1799998) has been reported for disease association studies [20, 21].

## 5 Conclusion

This study identified 29 deleterious CYP11B2 mutations using four computational tools including SIFT, Polyphen-2, I-Mutant, and Consurf. Then the structural analysis was performed using Sybyl and MD simulation to check the performance of Fadrozole over these mutations. The results have effectively shown that the fadrozole drug efficiently blocks to the identified damaging mutations thus is a very effective and potent drug.

## Acknowledgement

This project does not have any funding.

## Supporting Information Caption

S1 Table: 275 missense SNPs of CYP11B2

S2 Table: SIFT predicted Damaging SNPs

S3 Table: Polyphen-2 predicted damaging SNPs

S4 Table: I-Mutant predicted Damaging SNPs

S5 Table: Damaging nsSNPs by SIFT, POLYPHEN, I-Mutant and ConSurf

S6 Table: ConSurf predicted Conserved areas in 4FDHstructure of CYP11B2

## References

1. Karki R, Pandya D, Elston RC, Ferlini C. Defining “mutation” and “polymorphism” in the era of personal genomics. BMC Med Genomics. 2015;8(37):015–0115

2. Shastry BS. SNP alleles in human disease and evolution. J Hum Genet. 2002;47(11):561–6

3. Chen X, Sullivan PF. Single nucleotide polymorphism genotyping: biochemistry, protocol, cost and throughput. Pharmacogenomics J. 2003;3(2):77–96

4. Vaser R, Adusumalli S, Leng SN, Sikic M, Ng PC. SIFT missense predictions for genomes. Nat Protoc. 2016;11(1):1–9

5. Taymans SE, Pack S, Pak E, Torpy DJ, Zhuang Z, Stratakis CA. Human CYP11B2 (aldosterone synthase) maps to chromosome 8q24. 3. The Journal of Clinical Endocrinology & Metabolism. 1998;83(3):1033–6

6. Nanba K, Vaidya A, Williams GH, Zheng I, Else T, Rainey WE. Age-Related Autonomous Aldosteronism. Circulation. 2017;136(4):347–55

7. Feola M, Monteverde M, Vivenza D, Testa M, Leto L, Astesana V, et al. Prognostic Value of Different Allelic Polymorphism of Aldosterone Synthase Receptor in a Congestive Heart Failure European Continental Ancestry Population. Arch Med Res. 2017;48(2):156–61

8. Struthers AD, MacDonald TM. Review of aldosterone-and angiotensin II-induced target organ damage and prevention. Cardiovascular research. 2004;61(4):663–70

9. Jia M, Yang B, Li Z, Shen H, Song X, Gu W. Computational analysis of functional single nucleotide polymorphisms associated with the CYP11B2 gene. PloS one. 2014;9(8):e104311

10. Sherry ST, Ward MH, Kholodov M, Baker J, Phan L, Smigielski EM, et al. dbSNP: the NCBI database of genetic variation. Nucleic Acids Res. 2001;29(1):308–11

11. Kumar P, Henikoff S, Ng PC. Predicting the effects of coding non-synonymous variants on protein function using the SIFT algorithm. Nature protocols. 2009;4(7):1073

12. Ng PC, Henikoff S. Predicting deleterious amino acid substitutions. Genome research. 2001;11(5):863–74

13. Adzhubei IA, Schmidt S, Peshkin L, Ramensky VE, Gerasimova A, Bork P, et al. A method and server for predicting damaging missense mutations: Nat Methods. 2010 Apr;7(4):248–9. doi: 10.1038/nmeth0410-248.

14. Capriotti E, Calabrese R, Casadio R. Predicting the insurgence of human genetic diseases associated to single point protein mutations with support vector machines and evolutionary information2006.

15. Glaser F, Pupko T, Paz I, Bell RE, Bechor-Shental D, Martz E, et al. ConSurf: identification of functional regions in proteins by surface-mapping of phylogenetic information. Bioinformatics. 2003;19(1):163–4

16. Sybyl X. version 1.3. Tripos, LP: St Louis. 2011

17. Kollman PA, Massova I, Reyes C, Kuhn B, Huo S, Chong L, et al. Calculating structures and free energies of complex molecules: combining molecular mechanics and continuum models. Accounts of chemical research. 2000;33(12):889–97

18. Zhu M, Zhao S. Candidate gene identification approach: progress and challenges. International journal of biological sciences. 2007;3(7):420–7

19. Mooney SD, Klein TE. The functional importance of disease-associated mutation. BMC bioinformatics. 2002;3:24-.10.1186/1471-2105-3-24

20. Androulakis E, Tousoulis D, Papageorgiou N, Miliou A, Chatzistamatiou E, Moustakas G, et al. Effects of the C-344T aldosterone synthase gene variant on preclinical vascular alterations in essential hypertension. International journal of cardiology. 2013;168(2):1605–6

21. Freel E, Ingram M, Wallace A, White A, Fraser R, Davies E, et al. Effect of variation in CYP11B1 and CYP11B2 on corticosteroid phenotype and hypothalamic–pituitary–adrenal axis activity in hypertensive and normotensive subjects. Clinical endocrinology. 2008;68(5):700–6

